# Self-driving laboratories to autonomously navigate the protein fitness landscape

**DOI:** 10.1101/2023.05.20.541582

**Authors:** Jacob T. Rapp, Bennett J. Bremer, Philip A. Romero

## Abstract

Protein engineering has nearly limitless applications across chemistry, energy, and medicine, but creating new proteins with improved or novel functions remains slow, labor-intensive, and inefficient. In this work, we present the *Self-driving Autonomous Machines for Protein Landscape Exploration* (SAMPLE) platform for fully autonomous protein engineering. SAMPLE is driven by an intelligent agent that learns protein sequence-function relationships, designs new proteins, and sends designs to a fully automated robotic system that experimentally tests designed proteins and provides feedback to improve the agent’s understanding of the system. We deployed four SAMPLE agents with the goal of engineering glycoside hydrolase enzymes with enhanced thermal tolerance. Despite showing individual differences in their search behavior, all four agents quickly converged on thermostable enzymes that were at least 12 °C more stable than the starting sequences. Self-driving laboratories automate and accelerate the scientific discovery process and hold great potential for the fields of protein engineering and synthetic biology.

## Introduction

Human researchers engineer biological systems through the discovery-driven process of hypothesis generation, designing experiments to test hypotheses, performing these experiments in a wet laboratory, and interpreting the resulting data to refine understanding of the system. This process is iterated to converge on knowledge of biological mechanisms and design new systems with improved properties and behaviors. However, despite notable achievements in biological engineering and synthetic biology, this process remains highly inefficient, repetitive, and laborious, requiring multiple cycles of hypothesis generation and testing that can take years to complete.

Robot scientists and self-driving laboratories combine automated learning, reasoning, and experimentation to accelerate scientific discovery and design new molecules, materials, and systems. Intelligent robotic systems are superior to humans in their ability to learn across disparate data sources and data modalities, make decisions under uncertainty, operate continuously without breaks, and generate highly reproducible data with full metadata tracking and real-time data sharing. Autonomous and semi-autonomous systems have been applied to gene identification in yeast^1–3^, new chemical synthesis methodologies^4–6^, and the discovery of new photocatalysts^7^, photovoltaics^8^, adhesive materials^9^, and thin-film materials^10^. Self-driving laboratories hold great promise for the fields of protein engineering and synthetic biology^11–13^, but these applications are challenging because biological phenotypes are complex and nonlinear, genomic search spaces are high-dimensional, and biological experiments require multiple hands-on processing steps that are error prone and difficult to automate. There are examples of automated workflows for synthetic biology that require some human input and manual sample processing^14,15^, but are not fully autonomous in their ability to operate without human intervention.

We introduce the *Self-driving Autonomous Machines for Protein Landscape Exploration* (SAMPLE) platform to rapidly engineer proteins without human intervention, feedback, or subjectivity. SAMPLE is driven by an intelligent agent that learns protein sequence-function relationships from data and designs new proteins to test hypotheses. The agent interacts with the physical world though a fully automated robotic system that experimentally tests designed proteins by synthesizing genes, expressing proteins, and performing biochemical measurements of enzyme activity. Seamless integration between the intelligent agent and experimental automation enables fully autonomous design-test-learn cycles to understand and optimize the sequence-function landscape.

We deployed four independent SAMPLE agents to navigate the glycoside hydrolase landscape and discover enzymes with enhanced thermal tolerance. The agents’ optimization trajectories started with exploratory behavior to understand the broad landscape structure and then quickly converged on highly stable enzymes that were at least 12 °C more stable than the initial starting sequences. We observed notable differences in the individual agent’s search behavior arising from experimental measurement noise, yet all agents robustly identified thermostable designs while searching less than 2% of the full landscape. SAMPLE agents continually refine their understanding of the landscape through active information acquisition to efficiently discover optimized proteins. SAMPLE is a general-purpose protein engineering platform that can be broadly applied across biological engineering and synthetic biology.

## Results

### A fully autonomous system for protein engineering

We sought to build a fully autonomous system to mimic the human biological discovery and design process. Human researchers can be viewed as intelligent agents that perform actions in a laboratory environment and receive data as feedback. Through repeated interactions with the laboratory environment, human agents develop an understanding of the system and learn behaviors to achieve an engineering goal. SAMPLE consists of an intelligent agent that autonomously learns, makes decisions, and takes actions in a laboratory environment to explore protein sequence-function relationships and engineer proteins (**Fig 1a**).

**Figure 1:**
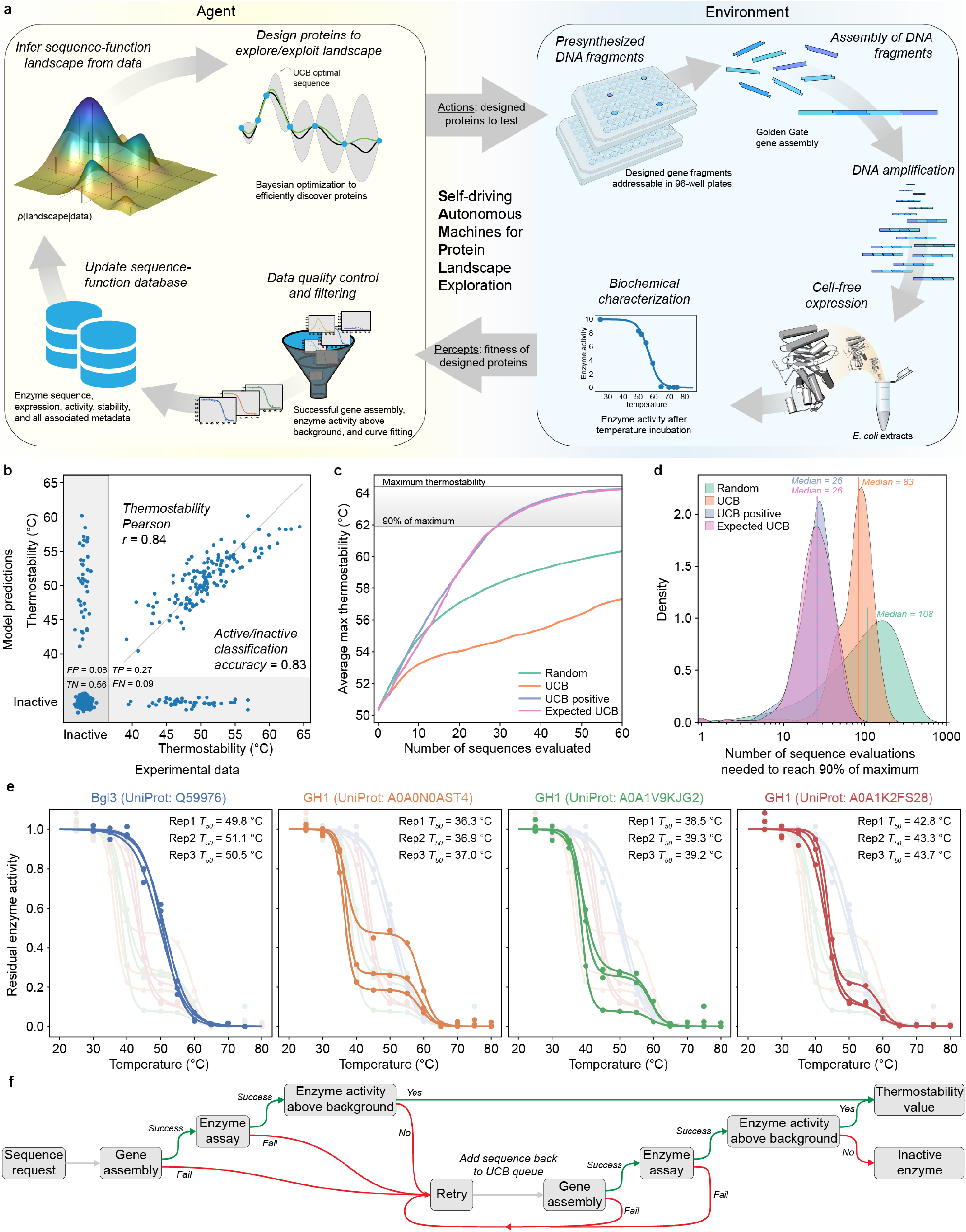
SAMPLE is a fully autonomous system for protein engineering. (**a**) SAMPLE consists of an intelligent agent that learns sequence-function relationships and designs proteins to test hypotheses. The agent sends designed proteins to a laboratory environment that performs fully automated gene assembly, protein expression, and biochemical characterization, and sends the resulting data back to the agent, which refines its understanding of the system and repeats the process. (**b**) The multi-output GP model classifies active/inactive P450s with 83% accuracy and predicts P450 thermostability with *r* = 0.84 using 10-fold cross-validation. (**c**) The performance of four sequential design strategies using P450 sequence-function data. The lines show the maximum observed thermostability for a given number of sequence evaluations, averaged over 10,000 simulated protein engineering trials. (**d**) The number of evaluations needed for the design strategies to discover sequences within 90% of the maximum thermostability (>61.9 °C) using 10,000 simulated protein engineering trials. (**e**) The reproducibility of the fully automated gene assembly, protein expression, and thermostability characterization pipeline on four diverse GH1 enzymes from Streptomyces species. The curves’ small shoulder centered around 60 °C is the result of background enzyme activity present in the *E. coli* cell extracts. (**f**) The pipeline has multiple layers of exception handing and data quality control for failed experimental steps.

The protein fitness landscape describes the mapping from sequence to function and can be imagined as a terrestrial landscape of peaks, valleys, and ridges^16^. The SAMPLE agent aims to identify high activity fitness peaks (i.e., top performing sequences) from an initially unknown sequence-function landscape. The agent actively queries the environment to gather information and construct an internal perception of the landscape. The agent must allocate resources between *exploration* to understand the landscape structure and *exploitation* that utilizes current landscape knowledge to identify optimal sequence configurations. We pose the agent’s protein engineering task as a Bayesian optimization (BO) problem that seeks to optimize an unknown objective function and must efficiently trade off between exploration and exploitation^17,18^.

The SAMPLE agent uses a Gaussian process (GP) model to build an understanding of the fitness landscape from limited experimental observations. The model must consider the protein function of interest, in addition to inactive “holes” in the landscape arising from destabilization of the protein structure^19,20^. We use a multi-output GP that simultaneously models whether a protein sequence is active/inactive and a continuous protein property of interest (See Supplementary Methods). We benchmarked our modeling approach on previously published cytochrome P450 data consisting of 331 inactive sequences and 187 active sequences with thermostability labels^21,22^. The multi-output GP showed excellent predictive ability with an 83% active/inactive classification accuracy and, for the subset of sequences that are active, predicts the thermostability with *r* = 0.84 (**Fig 1b**).

The GP model trained on sequence-function data represents the SAMPLE agent’s current knowledge, and from here, the agent must decide which sequences to evaluate next to achieve the protein engineering goal. BO techniques address this problem of sequential decision making under uncertainty. The upper-confidence bound (UCB) algorithm iteratively samples points with the largest upper-confidence bound (predictive mean plus prediction interval) and is proven to rapidly converge to the optimal point with high sample efficiency^23,24^. H^23,24^owever, naïve implementation of UCB for protein engineering is limited because the inactive “holes” in the landscape provide no information to improve the model. We devised two heuristic BO methods that consider the output of the active/inactive GP classifier (*p*_*active*_) to focus sampling toward functional sequences. The *UCB positive* method only considers the subset of sequences that are predicted to be active by the GP classifier (*p*_*active*_> 0.5) and selects the sequence with the top UCB value. The *Expected UCB* method takes the expected value of the UCB score by multiplying by the GP classifier *p*_*active*_ and selects the sequence with the top expected UCB value. We tested these methods by running 10,000 simulated protein engineering experiments with the cytochrome P450 data (**Fig 1cd**). On average, the UCB positive and Expected UCB methods found thermostable P450s with only 26 measurements and required 3-4 fold less samples than standard UCB and random. We also tested the BO methods in a batch setting where multiple sequences are tested in parallel and found a slight benefit to running experiments in smaller batches (**Fig S1**).

The agent designs proteins and sends them to the SAMPLE laboratory environment to provide experimental feedback. We developed a highly streamlined, robust, and general pipeline for automated gene assembly, cell-free protein expression, and biochemical characterization. Our procedure assembles pre-synthesized DNA fragments using Golden Gate cloning^25^ to produce a full intact gene and the necessary 5’/3’ untranslated regions for T7-based protein expression. The assembled expression cassette is then amplified via PCR and the product is verified using the fluorescent dye EvaGreen to detect double stranded DNA (**Fig S2**). The amplified expression cassette is then added directly to T7-based cell-free protein expression reagents to produce the target protein. Finally, the expressed protein is characterized using colorimetric/fluorescent assays to evaluate its biochemical activity and properties (**Fig S3**).

For this work, we focused on glycoside hydrolase enzymes and their tolerance to elevated temperatures. We tested the reproducibility of our automated experimental pipeline on four diverse glycoside hydrolase family 1 (GH1) enzymes from Streptomyces species (**Fig 1e**). The system reliably measured the thermostability (*T*_*50*_) of the enzymes with an error of less than 0.4 °C. The procedure takes approximately one hour for gene assembly, one hour for PCR, three hours for protein expression, three hours to measure thermostability, and overall nine hours to go from a requested protein design to a physical protein sample to a corresponding data point.

We added multiple layers of exception handling and data quality control to further increase the reliability of the SAMPLE platform (**Fig 1f**). The system checks whether (1) the gene assembly and PCR worked by assaying double stranded DNA with EvaGreen, (2) the enzyme reaction progress curves look as expected and the activity as a function of temperature can be fit using a sigmoid function, and (3) the observed enzyme activity is above the background hydrolase activity from the cell-free extracts. Failure at of any one of these checkpoints will flag the experiment as inconclusive and add the sequence back to the potential experiment queue.

### Designing combinatorial sequence spaces to broadly sample the protein landscape

The SAMPLE platform searches a large and diverse protein sequence space by assembling unique combinations of pre-synthesized DNA fragments. Combinatorial sequence spaces leverage exponential scaling to broadly sample the protein fitness landscape from a limited set of gene fragments. We define a combinatorial sequence space using a DNA assembly graph that specifies which sequence elements can be joined to generate a valid gene sequence (**Fig 2a**). We designed a GH1 combinatorial sequence space composed of sequence elements from natural GH1 family members, elements designed using Rosetta^26^, and elements designed using evolutionary information^27^. All designed sequence fragments are provided in **Table S1**. The full combinatorial sequence space contains 1352 unique GH1 sequences that differ by 116 mutations on average and by at least 16 mutations (**Fig 2b**). The sequences introduce diversity throughout the GH1 TIM barrel fold and sample up to six unique amino acids at each site (**Fig 2c**).

**Figure 2:**
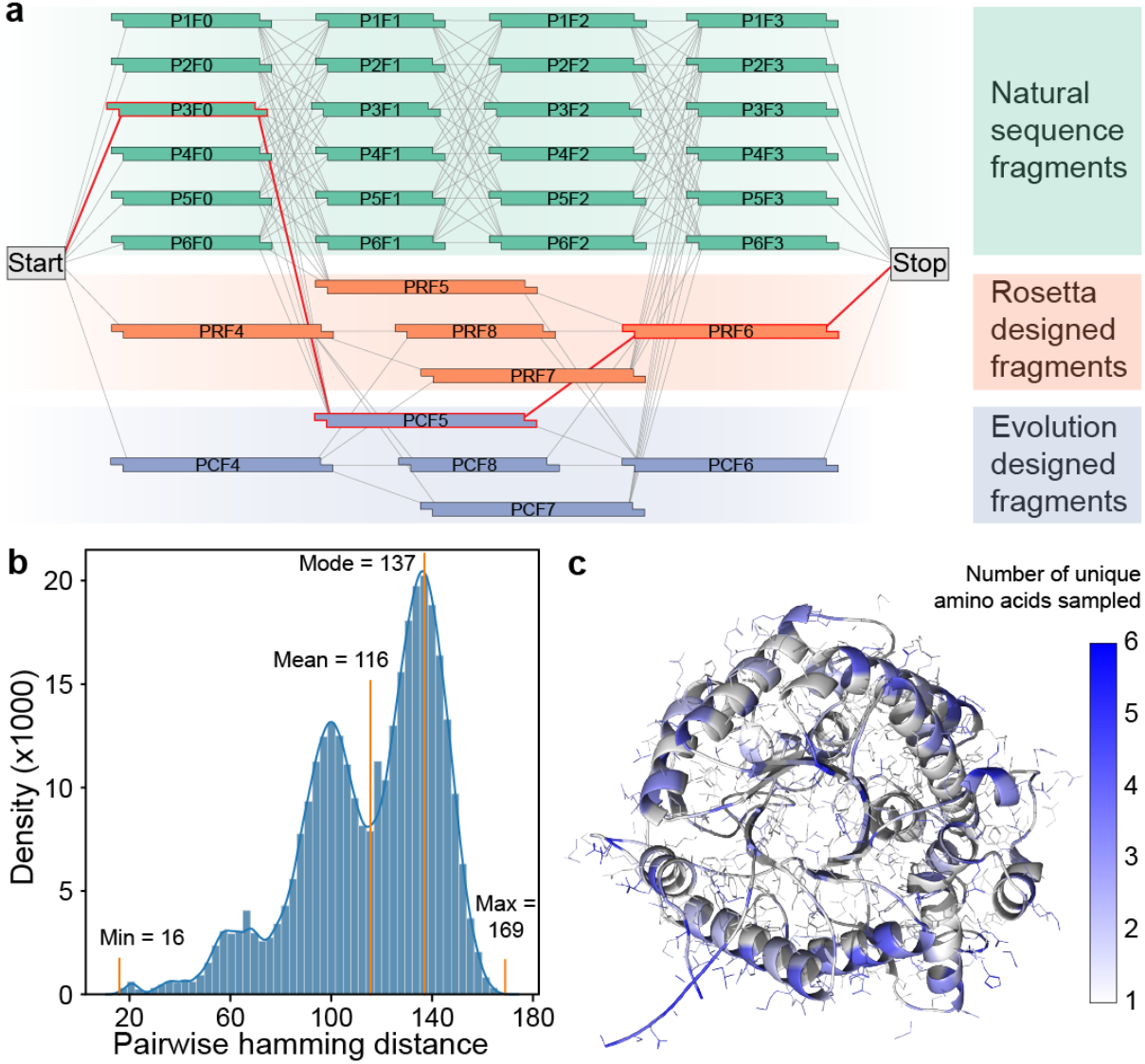
GH1 combinatorial sequence space. (**a**) A DNA assembly graph defines which sequence elements have compatible overhangs and can be joined to produce a valid gene sequence. Any path from the Start codon to the Stop codon (e.g., the red line) is a full gene sequence that can be assembled using Golden Gate cloning. Our GH1 sequence space has a total of 1352 paths from Start to Stop representing unique protein sequences. (**b**) Sequences within the designed GH1 sequence space differ by 116 amino acid substitutions on average and by at least 16 amino acids. (**c**) Sequences within this space sample amino acid diversity across the protein structure.

### Autonomous design of glycoside hydrolases via cloud-based laboratory environments

We applied SAMPLE with the goal of navigating and optimizing the GH1 thermostability landscape. We implemented our experimental pipeline on the Strateos Cloud Lab for enhanced scalability and accessibility by other researchers^28^. We deployed four independent SAMPLE agents that were each seeded with the same six natural GH1 sequences. The agents designed sequences according to the *Expected UCB* criterion, chose three sequences per round, and ran for a total of 20 rounds (**Fig 3a**). The four agents’ optimization trajectories showed a gradual climb of the landscape, with early phases characterized by exploratory behavior and later rounds consistently sampling thermostable designs. There were two instances where the quality filters missed faulty data and incorrectly assigned a thermostability value to an inactive sequence (Agent 1 in round 10 and Agent 3 in round 5). We intentionally did not correct these erroneous data points to observe how the agents recover from the error as they acquire more landscape information.

**Figure 3:**
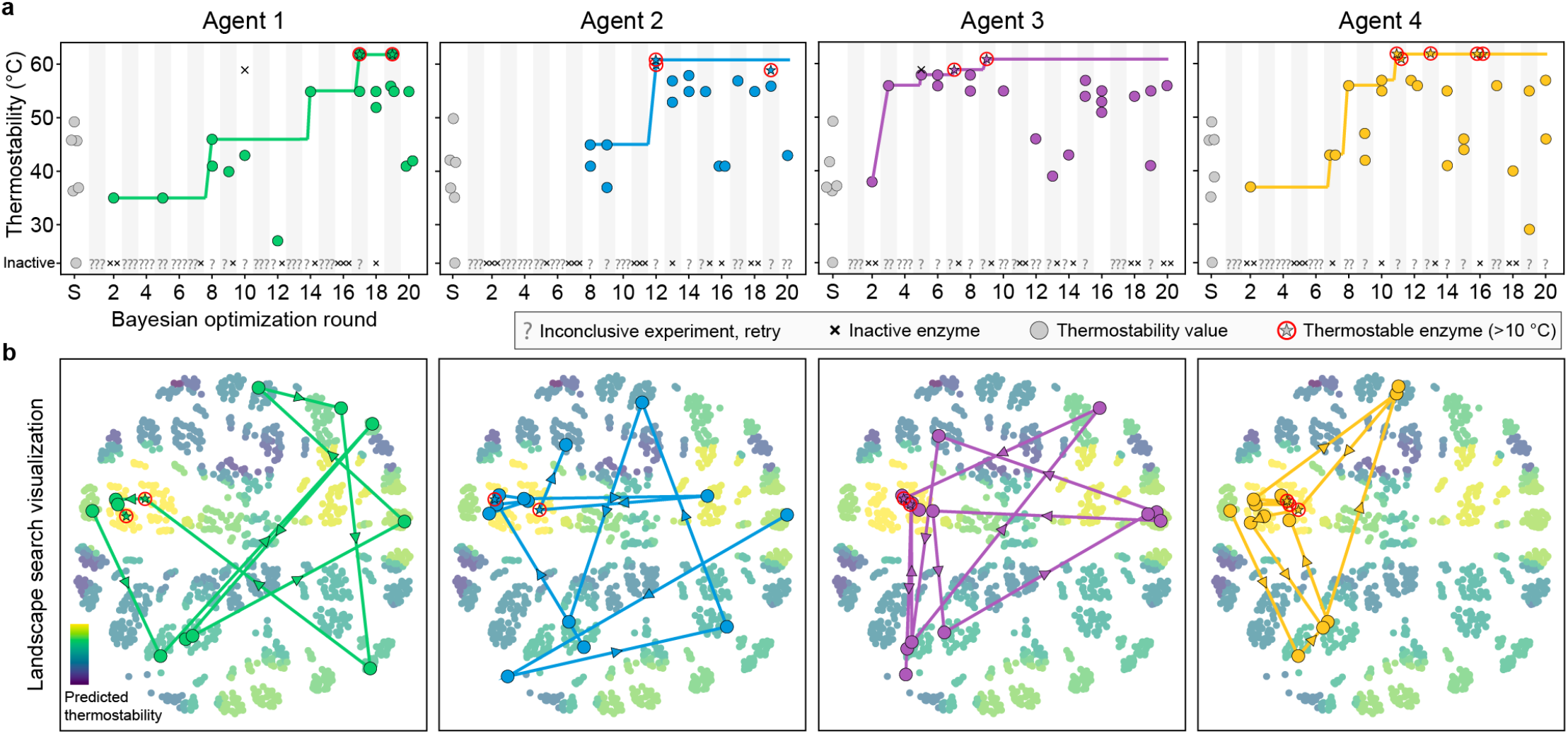
Autonomous exploration of the GH1 landscape. (a) Protein optimization trajectories of four independent SAMPLE agents. Inconclusive experiments, as defined from **Fig 1f**, are marked with a ‘?.’ There were two instances of inactive sequences that were incorrectly classified as active enzymes with thermostability values (Agent 1 in round 10 and Agent 3 in round 5). (b) Visualization of the landscape search. The 1352 possible sequences were arranged using multi-dimensional scaling and colored according to their predicted thermostability from the unified landscape model. The center left yellow cluster corresponds to the landscape’s fitness peak. The search trajectory is plotted as the most stable sequence from each round.

Each agent discovered GH1 sequences that were at least 12 °C more stable than the six initial natural sequences. The agents identify these sequences while searching less than 2% of the full combinatorial landscape. We visualized the agents’ search trajectory and found each agent broadly explored sequence space before converging on the same global fitness peak (**Fig 3b**). All four agents arrived upon similar regions of the landscape, but the top sequence discovered by each agent was unique. The thermostable sequences tended to be composed of the P6F0, P1F2 or P5F2, and P1F3 gene fragments, suggesting the corresponding amino acid segments may contain stabilizing residues and/or interactions.

The agents’ search trajectory and landscape ascent varied substantially despite being seeded with the same six sequences and following identical optimization procedures. Agent 3 found thermostable sequences by round 7, while Agent 1 took 17 rounds to identify similarly stable sequences. Agent 2 didn’t discover any functional sequences until round 8. The divergence in behaviors can be traced to the first decision-making step, where the four agents designed different sequences to test in round 1. These initial differences arose due to experimental noise in characterizing the six seed sequences that gave rise to slightly different landscape models that altered each agent’s subsequent decisions. The stochastic deviation between agents propagated further over the rounds to produce highly varied landscape searches, but that were ultimately steered back to the same global fitness peak.

### SAMPLE agents actively acquire information to explore and exploit protein fitness landscapes

SAMPLE agents efficiently and robustly discovered thermostable GH1 enzymes. We analyzed the four agents’ internal landscape perception and decision-making behavior to reveal how they navigate the protein fitness landscape. We plotted each agent’s model predictions for all 1352 combinatorial sequences over the course of the optimization (**Fig 4a**). The agents’ perception of the landscape changed over time and significant events, such as observing new stable sequences or erroneous data points, resulted in large landscape reorganization as indicated by crossing lines in **Fig 4a**. Many eventual top sequences were ranked near the bottom in early rounds.

**Figure 4:**
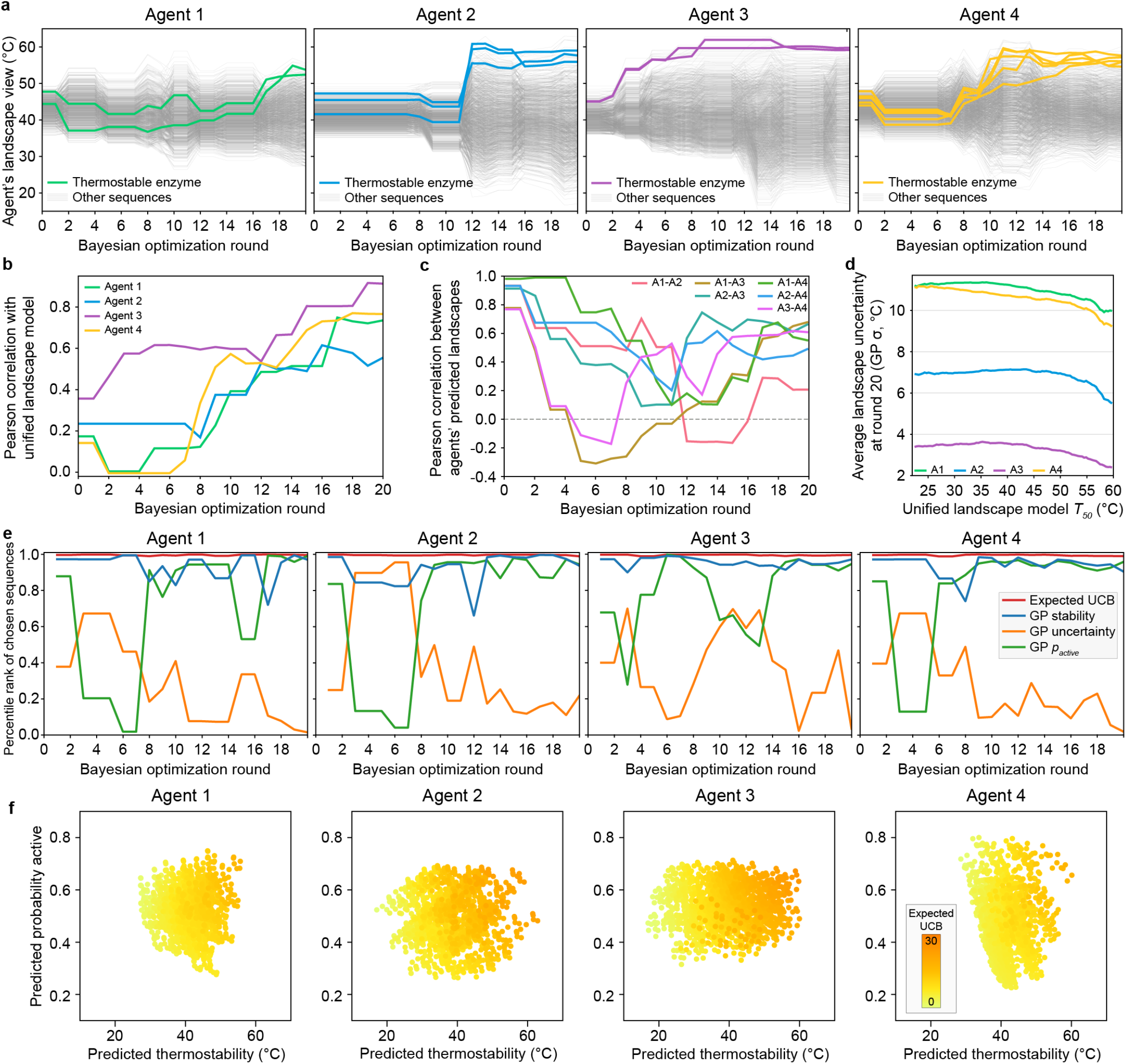
SAMPLE agents’ landscape search behavior. **(a)** Agents’ landscape perception over the course of the optimization. The light grey lines show the agent’s thermostability predictions for all 1352 sequences and the bold colored lines show sequences that were ultimately discovered to be thermostable by each agent. **(b)** Pearson correlation between the agents’ predicted thermostability landscape and the unified landscape model that incorporates all data retrospectively. **(c)** Pearson correlation between different agents’ thermostability landscapes over course of the optimization. **(d)** Average model uncertainty as a function of the landscape thermostability. The GP uncertainty (sigma) was averaged over all sequences falling within a 10 °C sliding window across the full *T*_*50*_ range predicted by the unified landscape model. **(e)** The chosen sequences’ percentile ranks for four key factors including Expected UCB, the thermostability model’s mean prediction (GP stability), the thermostability model’s predictive uncertainty (GP uncertainty), and the active/inactive classifier’s predicted probability a sequence is active (*p*_*active*_). The percentile ranks were averaged over the three sequences in the batch. A percentile rank approaching one indicates the chosen sequences were exceptional for a given factor. **(f)** The agent’s view of the landscape after the 20 rounds of optimization with expected UCB overlaid to highlight which factors are contributing to expected UCB.

To get an estimated “ground truth” landscape, we trained a GP model on all sequence-function data from all agents, which we refer to as the *unified landscape model* (**Fig S4**). We analyzed how each agent’s landscape perception correlates with the unified landscape model and found agents’ understanding became progressively refined and improved as they acquired sequence-function information (**Fig 4b**). Notably, most agents discovered thermostable sequences by rounds 11-12, when their understanding of the landscape was still incomplete, as indicated by a moderate Pearson correlation of ∼0.5. We also analyzed the different agents’ degree of agreement on the underlying landscape structure (**Fig 4c**). All four agents started with correlated landscape perceptions because they were initialized from the same six sequences, but the landscape consistency quickly dropped, with some agents even displaying negative correlations. The early disagreement arose because each agent pursued a unique search trajectory and thus specialized on different regions of the landscape. The correlation between agents’ perceived landscapes eventually increased as more information was acquired. Again, it’s notable how the agents tended to discover thermostable sequences by rounds 11-12, while largely disagreeing on the full landscape structure. BO algorithms are efficient because they focus on understanding the fitness peaks, while devoting less effort to regions known to be suboptimal. After round 20, we found the four agents were more confident on the top thermostable sequences and had greater uncertainty associated with lower fitness regions of the landscape (**Fig 4d**).

The SAMPLE agents designed sequences according to the *Expected UCB* criterion, which considers the thermostability prediction, the model uncertainty, and the probability an enzyme is active (*p*_*active*_). We wanted to understand the interplay of these three factors and how they influenced each agent’s decision making. We looked at the sequences chosen each round and their percentile rank for thermostability prediction, model uncertainty, and *p*_*active*_ (**Fig 4e**). The agents prioritized the thermostability prediction throughout the optimization, and tended to sample uncertain sequences in early phases, while emphasizing *p*_*active*_ in the later phases. Agent 3 prioritized *p*_*active*_ earlier than the other agents, which seems to be the result of discovering thermostable sequences early and putting less emphasis on exploration. We also analyzed the agents’ final perception of thermostability, *p*_*active*_, and expected UCB and found the agents specialized on different factors resulting from their past experiences (**Fig 4f**). Agent 4’s expected UCB is dictated by its large *p*_*active*_ range, while Agent 2’s is determined by its predicted thermostability. Meanwhile, Agent 3 still has considerable landscape uncertainty as indicated by the high expected UCB points with moderate thermostability and *p*_*active*_ predictions.

### Human validation and characterization of machine-designed proteins

The SAMPLE system was given a protein engineering objective, reagents, and DNA components, and autonomously proceeded to search the fitness landscape and discover thermostable GH1 enzymes. We experimentally characterized the top sequence discovered by each agent to validate the SAMPLE system’s findings. We found all four machine-designed enzymes were significantly more thermostable than the top natural sequence (Bgl3) and the designs from Agents 1 and 4 were nearly 10 °C more stable (**Fig 5a**). The thermostability differences were not as large as observed using our automated experimental setup and this is likely a result of different protein expression and assay conditions. We also tested the enzymes’ kinetic properties and found all designs displayed Michaelis-Menten kinetics with catalytic efficiencies (*k*_*cat*_/*K*_*M*_) that match or exceed wild-type Bgl3 (**Fig 4b**). Our protein engineering objective did not explicitly seek to find more catalytically efficient enzymes, but the search may have been biased toward more active enzymes due to data filtering steps that excluded enzymes with low activity.

**Figure 5:**
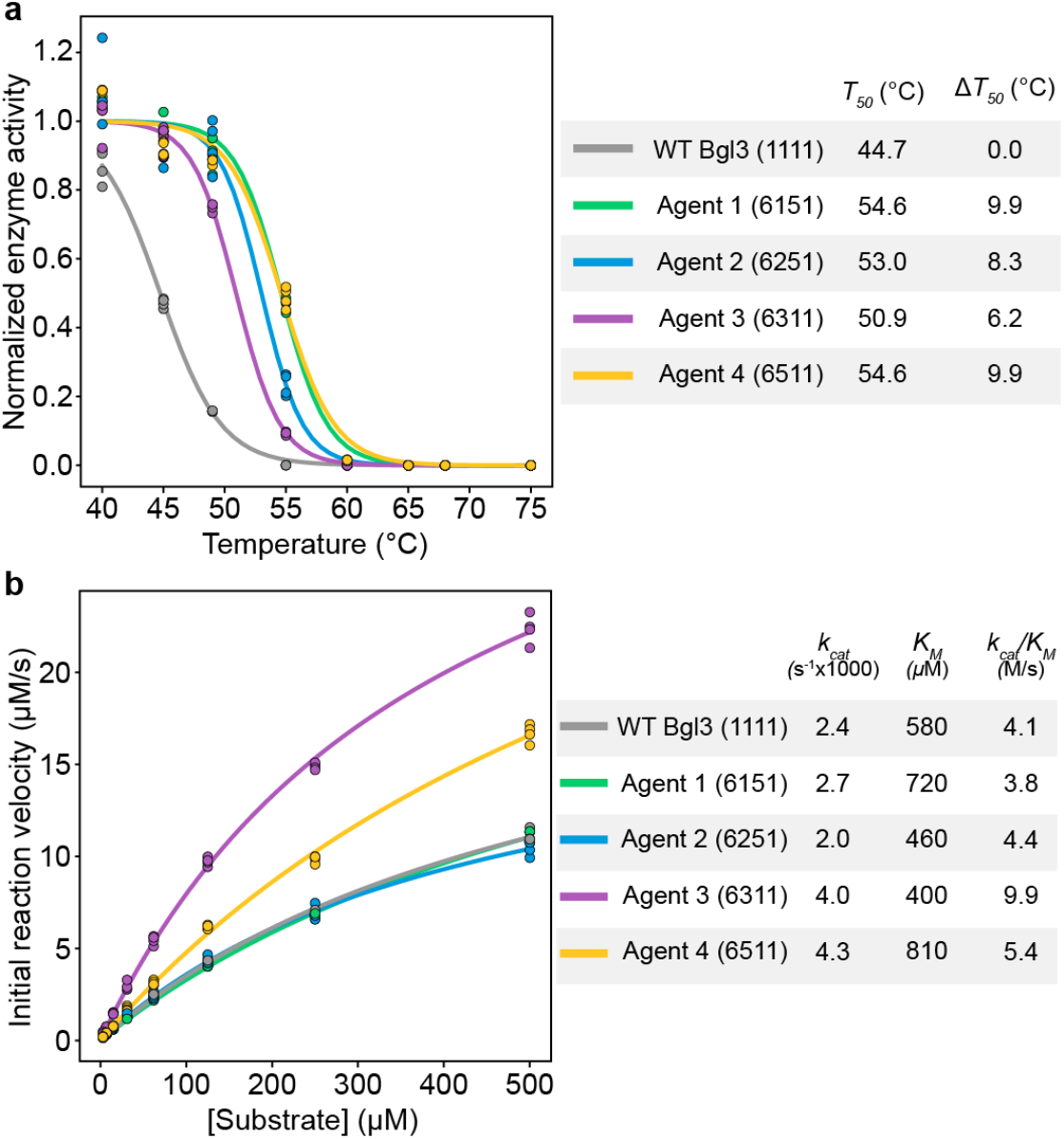
Thermostability and kinetic properties designed GH1s. **(a)** Enzyme inactivation as a function of temperature. Each measurement was performed in quadruplicate and shifted sigmoid functions were fit to the average over replicates. The *T*_*50*_ parameter is the midpoint of the sigmoid function and is defined as the temperature where 50% of the enzyme is irreversibly inactivated in 10 minutes. The enzyme variant is specified by the sequence of its four constituent fragments, e.g. 6151 corresponds to P6F0-P1F1-P5F2-P1F3. **(b)** Enzyme reaction velocity as a function of substrate concentration. Each measurement was performed in triplicate and the Michaelis-Menten equation was fit to the average over replicates to determine the kinetic constants.

## Discussion

Self-driving laboratories automate and accelerate the scientific discovery process and hold great potential to revolutionize the fields of protein engineering and synthetic biology. Automating the biological design process remains challenging due to the scale and complexity of biological fitness landscapes and the specialized operations required for wet lab experiments. In this work, we developed the *Self-driving Autonomous Machines for Protein Landscape Exploration* (SAMPLE) platform for fully autonomous protein engineering. SAMPLE tightly integrates automated learning, decision making, protein design, and experimentation to explore fitness landscapes and discover optimized proteins. We deployed SAMPLE agents with the goal of engineering glycoside hydrolase (GH1) enzymes with enhanced thermal tolerance. The agents efficiently and robustly searched the landscape to identify thermostable enzymes that were at least 12 °C more stable than the initial starting sequences.

SAMPLE is a general protein engineering platform that can be readily applied to diverse protein engineering targets. The system does not require prior knowledge of protein structure or mechanism, but instead takes an unbiased approach that examines how sequence changes impact the target function. The greatest barrier to establishing SAMPLE for a new protein target is the required biochemical assay. The robotic systems used in this work had access to a microplate reader and thus required a colorimetric or fluorescence-based assay. In principle, more advanced analytical instruments, such as liquid chromatography-mass spectrometry (LC-MS) or nuclear magnetic resonance (NMR) spectroscopy, could be integrated into automation systems to expand the types of protein functions that could be engineered. Finally, we implemented our full experimental pipeline on the Strateos Cloud Lab to produce a cost effective and accessible system that can be adopted by other synthetic biology researchers.

SAMPLE has potential to streamline and accelerate the process of protein engineering. The experimental side of the system is the major throughput bottleneck that limits the overall process. A single round of experimental testing takes 9 hours on our Tecan automation system or 10 hours split over two days (5 hrs x 2 days) on the Strateos Cloud Lab. At these rates, with continuous operation, the system could get through 20 design-test-learn cycles in just 1-2 weeks. In practice the process was much slower due to system downtime, robotic malfunctions, and time needed for restocking reagents. Our 20 rounds of GH1 optimization took just under six months, which included a single 2.5 month pause caused by shipping delays. Even this six-month duration compares favorably to human researchers, which we estimate may take 6-12 months to perform similar experiments using standard molecular biology and protein engineering workflows. Learning from previous delays, and with better planning, we estimate SAMPLE could perform 20 design-test-learn cycles in two months using the Strateos Cloud Lab.

We deployed four identical SAMPLE agents and observed notable differences in their search behavior and landscape optimization efficiency. The agents explored distinct regions of sequence space, specialized on different tasks such as classifying active/inactive enzymes versus predicting thermostability, and Agent 3 discovered thermostable enzymes with 10 fewer rounds than Agent 1. The initial divergence in behavior arises from experimental measurement noise, which influences the agents’ decisions, which then further propagates differences between agents. There is also an element of luck that is compounded with positive feedback: an agent may happen to search in a particular region and come across improved sequences, which then drives the search upward in favorable directions. These observations have interesting parallels with human researchers, where success or failure could be influenced by seemingly inconsequential experimental outcomes and the resulting decisions. The SAMPLE agents explored distinct regions of the landscape and specialized on unique tasks, which indicates potential to coordinate multiple agents towards a single protein engineering goal. The decentralized and on-demand nature of cloud laboratory environments would further assist multi-agent coordination systems.

Other research groups have developed automated pipelines and semi-autonomous systems for biological systems engineering. Carbonell et al developed an automated design-build-test-learn pipeline that searches over gene regulatory elements such as promoters and operon configurations to optimize biosynthetic pathway titers^14^. They demonstrated their pipeline by performing two design-build-test-learn cycles to optimize flavonoid and alkaloid production in *E. coli*. Each step of this pipeline utilized automation, but the entire procedure was not fully integrated to enable autonomous operation. HamediRad et al developed automated design-build-test-learn system to optimize biosynthetic pathways by searching over promoters and ribosome binding sites^15^. They applied their system to enhance lycopene production in *E. coli* and performed three design-build-test-learn cycles. The most notable difference between these prior demonstrations is SAMPLE’s high level of autonomy, which allowed us to perform four independent trials of 20 design-test-learn cycles each. High autonomy enables more experimental cycles without the need for slow human intervention.

The powerful combination of artificial intelligence and automation is disrupting nearly every industry from manufacturing and food preparation to pharmaceutical discovery, agriculture, and waste management. Self-driving laboratories will revolutionize the fields of biomolecular engineering and synthetic biology by automating highly inefficient, time consuming, and laborious protein engineering campaigns, enabling rapid turnaround, and allowing researchers to focus on important downstream applications. Intelligent autonomous systems for scientific discovery will become increasingly powerful with continued advances in deep learning, robotic automation, and high-throughput instrumentation.

## Supplementary Materials and Methods

### Benchmarking Bayesian optimization methods on cytochrome P450 data

We compiled a cytochrome P450 data set to benchmark modeling and Bayesian optimization methods. The data set consists of 518 data points with binary active/inactive data from Otey et al.^22^ and thermostability measurements from Li et al.^21^. We tested the multi-output GP model by performing 10-fold cross-validation where a GP classifier was trained on binary active/inactive data and a GP regression model was trained on thermostability data. The models used a linear Hamming kernel (sklearn^29^ DotProduct with sigma_0 = 1) with an additive noise term (sklearn WhiteKernel noise_level = 1). For the test set predictions, we categorized sequences as either true negative (TN), false negative (FN), false positive (FP), or true positive (TP), and for true positives we calculated the correlation between predicted thermostability and true thermostability values.

We used the cytochrome P450 data to benchmark the Bayesian optimization (BO) methods. The *Random* method randomly selects a sequence from the pool of untested sequences. The upper-confidence bound (*UCB*) method chooses the sequence with the largest upper-confidence bound (GP thermostability model mean + 95% prediction interval) from the pool of untested sequences. The *UCB* method does not have an active/inactive classifier and if it observes an inactive sequence, it does not update the GP regression model. The *UCB positive* method incorporates the active/inactive classifier and only considers the subset of sequences that are predicted to be active by the GP classifier (*p*_*active*_> 0.5). From this subset of sequences it selects the sequence with the top UCB (GP thermostability model mean + 95% prediction interval) value. The *Expected UCB* method takes the expected value of the UCB score by (1) subtracting the minimum value from all thermostability predictions to set the baseline to zero, (2) adding the 95% prediction interval, and (3) multiplying by the active/inactive classifier *p*_*active*_. The sequence with the top expected UCB value is chosen from the pool of untested sequences.

We tested the performance of these four methods by running 10,000 simulated protein engineering trials using the cytochrome P450 data. For each simulated protein engineering trial, the first sequence was chosen randomly, and subsequent experiments were chosen according to the different BO criteria. A trial’s performance at a given round is the maximum observed thermostability from that round and all prior rounds. We averaged each performance profile over the 10,000 simulated trials.

We also developed and tested batch methods that select multiple sequences each round. For the batch methods we use the same UCB variants described above to choose the first sequence in the batch, then we update the GP model assuming the chosen sequence is equal to its predicted mean, and then we select the second sequence according to the specified UCB criteria. We continue to select sequences and update the GP model until the target batch size is met. We assessed how the batch size affects performance by running 10,000 simulated protein engineering trials at different batch sizes and evaluating how many learning cycles were needed to reach 90% of the maximum thermostability.

### Glycoside hydrolase (GH1) combinatorial sequence space design

We designed a combinatorial glycoside hydrolase family 1 (GH1) sequence space composed of sequence elements from natural GH1 family members, elements designed using Rosetta^26^, and elements designed using evolutionary information^27^. The combinatorial sequence space mixes and matches these sequence elements to create new sequences. The sequences are assembled using Golden Gate cloning and thus require common four base pair overhangs to facilitate assembly between adjacent elements.

We choose six natural sequences by running a BLAST search on Bgl3^30^ and selecting five additional sequences that fell within the 70-80% sequence identity range (**Fig S3**). We aligned these six natural sequences and choose breakpoints using SCHEMA recombination^31,32^ with the wild-type Bgl3 crystal structure (PDB ID: 1GNX). The breakpoints for the Rosetta and evolution-designed sequence fragments were chosen to interface with the natural fragments and also introduce new breakpoints to promote further sequence diversity. For the Rosetta fragments, we started with the crystal structure of wild-type Bgl3 (PDB ID: 1GNX), relaxed the structure using FastRelax, and used RosettaDesign to design a sequence segment for a given fragment while leaving the remainder of the sequence and structure as wild-type Bgl3. At each position, we only allowed residues that were observed within the six aligned natural sequences. For the evolution designed fragments, we used Jackhmmer^33^ to build a large family multiple sequence alignment and designed sequence segments containing the most frequent amino acid from residues that were observed within the six natural sequences.

We designed DNA constructs to assemble sequences from the combinatorial sequence space using Golden Gate cloning. The designed amino acid sequence elements were reverse translated using the Twist codon optimization tool and the endpoints were fixed to preserve the correct Golden Gate overhangs. We added BsaI sites to both ends to allow restriction digestion and ordered the 34 gene fragments cloned into the pTwist Amp High Copy vector. Each sequence element’s amino acid and gene sequence are given in **Table S1**.

### Automated gene assembly, protein expression, and protein characterization

We implemented our fully automated protein testing pipeline on an in-house Tecan liquid handing system and the Strateos Cloud Lab. The system is initialized with a plate of the 34 gene fragments (5 ng/μL), NEB Golden Gate Assembly Kit (E1601L) diluted to a 2x stock solution, a 2 μM solution of forward and reverse PCR primers, Phusion 2X Master Mix (ThermoFisher F531L), 2x EvaGreen stock solution, Bioneer AccuRapid Cell Free Protein Expression Kit (Bioneer K-7260) Master Mix diluted in water to 0.66x, AccuRapid *E. coli* extract with added 40 μM fluorescein, a fluorogenic substrate master mix (139 μM 4-Methylumbelliferyl-α-D-Glucopyranoside, 0.139% v/v dimethyl sulfoxide, 11 mM phosphate, and 56 mM NaCl), and water.

#### Golden Gate assembly of DNA fragments

For a given assembly, 5 μL of each DNA fragment are mixed and 10 μL of the resultant mixture is then combined with 10 μL of 2x Golden Gate Assembly Kit. This reaction mix is heated at 37 °C for 1 hour, followed by a 5-minute inactivation at 55 °C.

#### PCR amplification of assembled genes

10 μL of the Golden Gate assembly product is combined with 90 μL of the PCR primers stock, and 10 μL of this mixture is then added to 10 μL Phusion 2X Master Mix. PCR was carried out with a 5-minute melt at 98 °C, followed by 35 cycles of 56 °C anneal for 30 seconds, 72 °C extension for 60 seconds, and 95 °C melt for 30 seconds. This is followed by one final extension for 5 minutes at 72 °C.

#### Verification of PCR amplification

10 μL of the PCR product is combined with 90 μL of water, and 50 μL of this mixture is then combined with 50 μL 2X EvaGreen. The fluorescence of the sample is read on a microplate reader (ex: 485 nm/em: 535 nm) and the signal is compared to prior positive/negative control PCRs to determine whether PCR amplification was successful.

#### Cell-free protein expression

30 μL of the 10x PCR dilution from the previous step are added to 40 μL AccuRapid E. coli extract and mixed with 80 μL the AccuRapid Master Mix. The protein expression reaction is incubated at 30 °C for 3 hours.

#### Thermostability assay

70 μL of the expressed protein is diluted with 600 μL water, and 70 μL aliquots of this diluted protein are added to a column of a 96-well PCR plate for temperature gradient heating. The plate is heated for 10 minutes on a gradient thermocycler such that each protein sample experiences a different incubation temperature. After incubation, 10 μL of the heated sample is added to 90 μL of the fluorogenic substrate master mix and mixed by pipetting. The fluorescein internal standard is analyzed on a microplate reader (ex: 494 nm/em: 512 nm) for sample normalization and then enzyme reaction progress is monitored by analyzing the sample fluorescence (ex: 372 nm/em: 445 nm) every 2 minutes for an hour.

### Human characterization of top designed enzymes

#### Bacterial protein expression and purification

The designs were built using Golden Gate cloning to assemble the constituent gene fragments and the full gene was cloned into the pET-22b expression plasmid. The assemblies were transformed into E. coli DH5α cells and the gene sequences were verified using Sanger sequencing. The plasmids were then transformed into E. coli BL21(DE3) and preserved as glycerol stocks at -80 °C. The glycerol stocks were used to inoculate an overnight Luria broth (LB) starter culture and the next day this culture was diluted 100x into a 50 mL LB expression culture with 50 μg/mL carbenicillin. The culture was incubated shaking at 37 °C until the OD600 of reached 0.5-0.6 and then induced with 1 mM IPTG. The expression cultures were incubated shaking overnight at 16 °C and the next day the cultures were harvested by centrifugation at 3600 g for 10 minutes and discarding the supernatant. The cell pellets were resuspended in 5 mL PBS and lysed by sonication at 22 W for 20 cycles of 5 seconds on and 15 seconds off. The lysates were clarified by centrifugation 10,000 g for 15 minutes.

The enzymes were purified by loading the clarified lysates onto a Ni-NTA agarose column (Cytiva 17531801), washing with 20 mL wash buffer (25 mM Tris, 400 mM NaCl, 20 mM Imidazole, 10% v/v glycerol, pH 7.5), and eluting with 5 mL elution buffer (25 mM Tris, 400 mM NaCl, 250 mM Imidazole, 10% v/v glycerol, pH 7.5). The eluted samples were concentrated using an Amicon filter concentrator and concurrently transitioned to storage buffer (25 mM Tris, 100 mM NaCl, 10% v/v glycerol, pH 7.5). The final protein concentration was determined using the Bio-Rad protein assay, the samples were diluted to 2 mg/mL in storage buffer, and frozen at -80 °C.

#### Thermostability assay

The clarified cell lysate from the protein expression was diluted 100x in PBS. 100 μL of the diluted lysate was arrayed into a 96-well PCR plate and heated for 10 minutes on a gradient thermocycler from 40 °C to 75 °C. The heated samples were assayed for enzyme activity in quadruplicate with final reaction conditions of 10% heated lysate, 125 μM 4-Methylumbelliferyl-β-D-Glucopyranoside, 0.125% v/v DMSO, 10 mM phosphate buffer pH 7, and 50 mM NaCl. The reaction progress was monitored using a microplate reader analyzing sample fluorescence (ex: 372 nm/em: 445 nm) every 2 minutes for 30 minutes. The reaction progress curves were fit using linear regression to get the reaction rate and a shifted sigmoid function was fit to the rate as a function of temperature incubation to obtain the *T*_*50*_ value.

#### Michaelis-Menten kinetic assay

The purified enzymes were assayed in quadruplicate along an eight-point twofold dilution series of 4-Methylumbelliferyl-β-D-Glucopyranoside starting from 500 μM. The assays were performed with 10 nM enzyme, 0.5% v/v DMSO, 10 mM phosphate buffer pH 7, and 50 mM NaCl. The reaction progress was monitored using a microplate reader analyzing sample fluorescence (ex: 372 nm/em: 445 nm) every 2 minutes for 30 minutes. The initial rate data was fit the Michaelis-Menten equation using the scikit-learn^29^ curve_fit function to determine the enzyme *k*_*cat*_ and *K*_*M*_.

## Supplementary Figures

**Figure S1:**
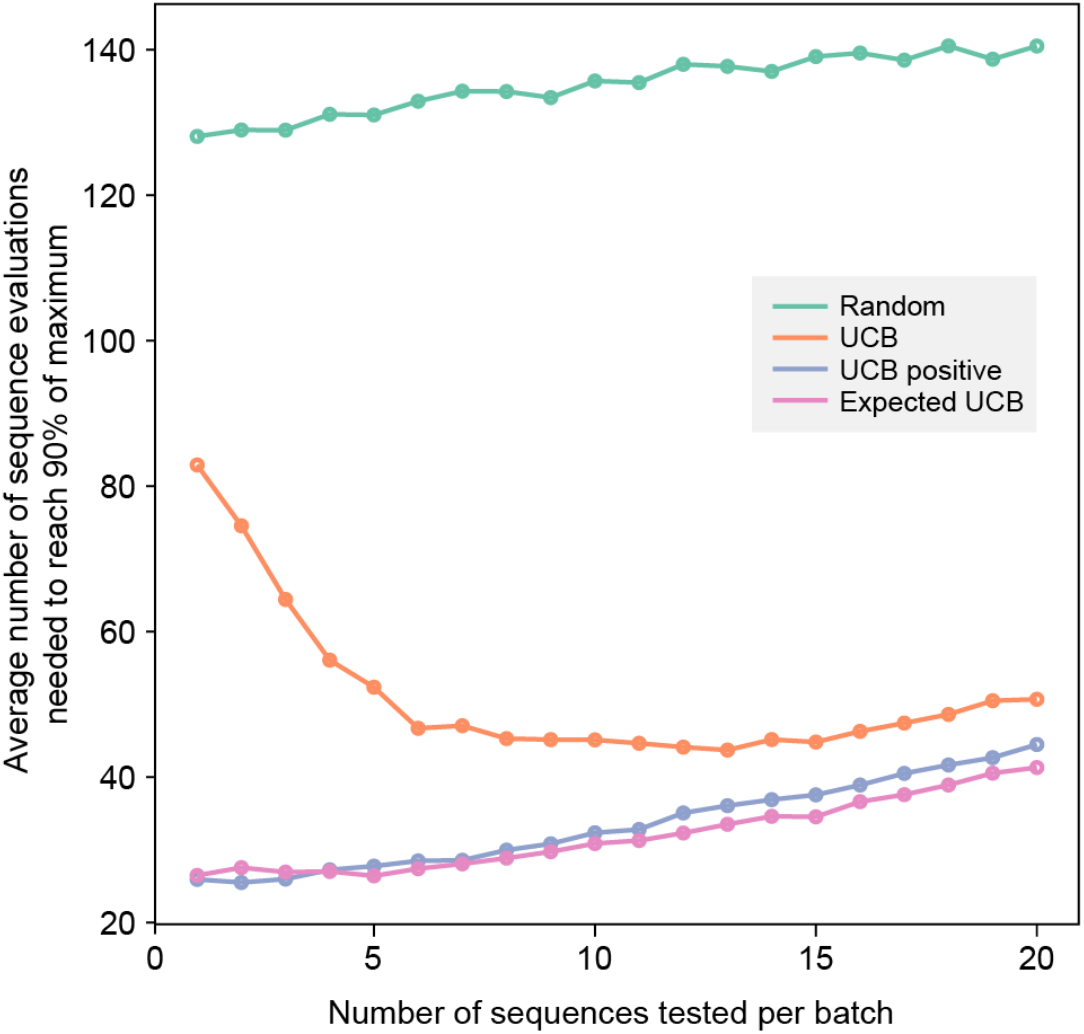
Batch BO method performance. We tested batch BO methods by running 10,000 simulated protein engineering experiments with the cytochrome P450 data and evaluating how many sequence evaluations were needed to discover sequences within 90% of the maximum sequence (>61.9 °C). The UCB positive and Expected UCB methods show the best overall performance and perform better with smaller batch sizes.

**Figure S2:**
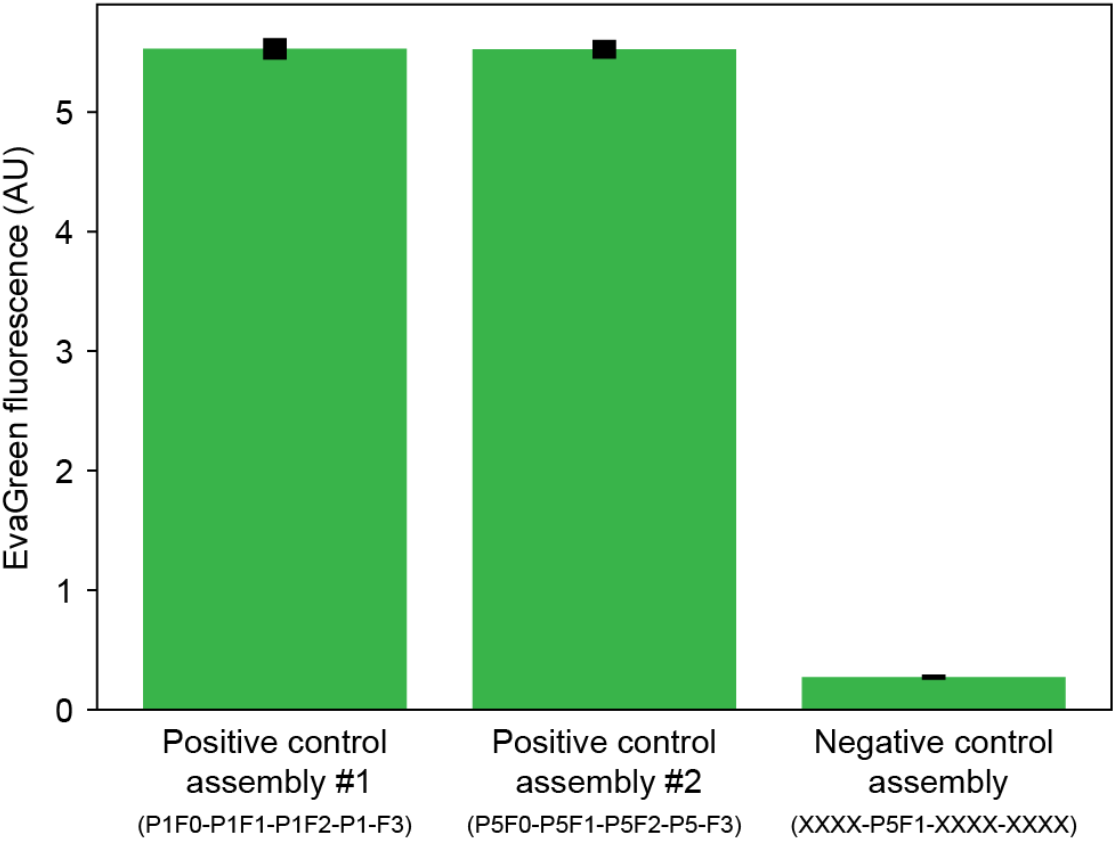
EvaGreen assay to test successful gene assembly and PCR amplification. We tested two positive control assemblies with DNA fragments to assemble full genes and a negative control assembly that only had one DNA fragment. The EvaGreen fluorescence clearly distinguishes between successful and unsuccessful gene assembly.

**Figure S3:**
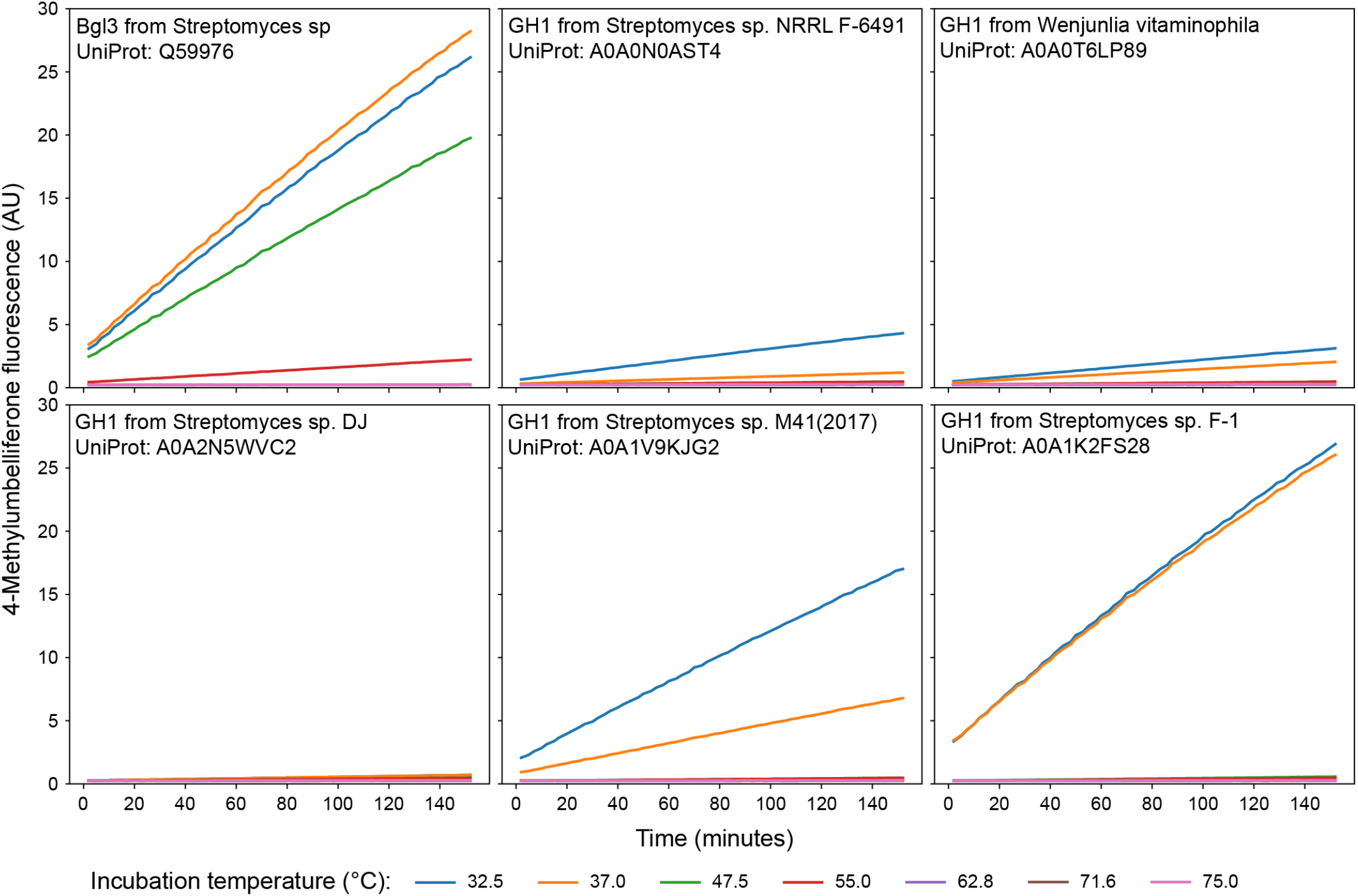
Enzyme reaction progress curves. An example of enzyme reaction progress curves for the six natural GH1s used to initialize the BO runs. Enzyme reactions were run at room temperature after a 10-minute incubation at the specified temperature. The decrease in activity is the result of irreversible enzyme inactivation from the temperature incubation.

**Figure S4:**
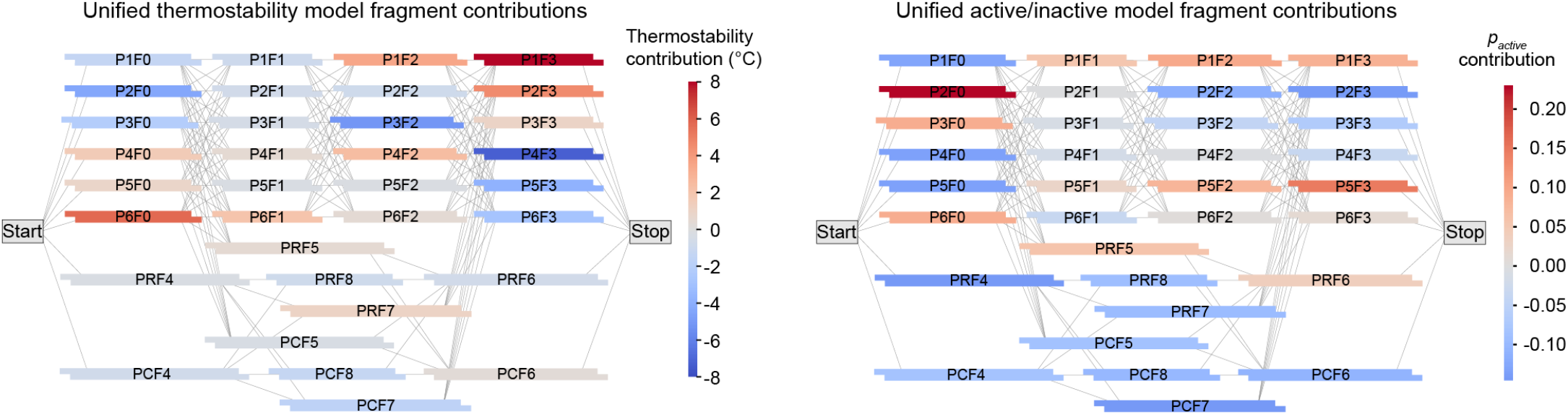
The contribution of individual gene fragments to enzyme thermostability and probability of being active. The unified landscape model was trained on all collected data across all agents and all runs. The fragment contributions are calculated as the mean of the property (thermostability or *p*_*active*_) across all sequences subtracted from the mean over sequences that have that specific fragment.

